# Simplified fecal community transplant restores *Clostridioides difficile* colonization resistance to antibiotic perturbed murine communities

**DOI:** 10.1101/2022.05.11.491589

**Authors:** Nicholas A. Lesniak, Sarah Tomkovich, Andrew Henry, Ana Taylor, Joanna Colovas, Lucas Bishop, Kathryn McBride, Patrick D. Schloss

## Abstract

Fecal communities transplanted into individuals can eliminate recurrent *Clostridioides difficile* infection (CDI) with high efficacy. However, this treatment is only used once CDI becomes resistant to antibiotics or has recurred multiple times. We sought to investigate whether a fecal community transplant (FCT) pre-treatment could be used to prevent CDI altogether. We treated male C57BL/6 mice with either clindamycin, cefoperazone, or streptomycin, and then inoculated them with the microbial community from untreated mice before challenging with *C. difficile*. We measured colonization and sequenced the V4 region of the 16S rRNA gene to understand the dynamics of the murine fecal community in response to the FCT and *C. difficile* challenge. Clindamycin-treated mice became colonized with *C. difficile* but cleared it naturally and did not benefit from the FCT. Cefoperazone-treated mice became colonized by *C. difficile*, but the FCT enabled clearance of *C. difficile*. In streptomycin-treated mice, the FCT was able to prevent *C. difficile* from colonizing. Then we diluted the FCT and repeated the experiments. Cefoperazone-treated mice no longer cleared *C. difficile*. However, streptomycin-treated mice colonized with 1:10^2^ dilutions resisted *C. difficile* colonization. Streptomycin-treated mice that received a FCT diluted 1:10^3^, *C. difficile* colonized but later was cleared. In streptomycin-treated mice, inhibition of *C. difficile* was associated with increased relative abundance of a group of bacteria related to *Porphyromonadaceae* and *Lachnospiraceae*. These data demonstrate that *C. difficile* colonization resistance can be restored to a susceptible community with a FCT as long as it complements the missing populations.

**Importance:** Antibiotic use, ubiquitous with the healthcare environment, is a major risk factor for *Clostridioides difficile* infection (CDI), the most common nosocomial infection. When *C. difficile* becomes resistant to antibiotics, a fecal microbiota transplant from a healthy individual can effectively restore the gut bacterial community and eliminate the infection. While this relationship between the gut bacteria and CDI is well established, there are no therapies to treat a perturbed gut community to prevent CDI. This study explored the potential of restoring colonization resistance to antibiotic-induced susceptible gut communities. We described the effect gut bacteria community variation has on the effectiveness of a fecal community transplant for inhibiting CDI. These data demonstrated that communities susceptible to CDI can be supplemented with fecal communities but the effectiveness depended on the structure of the community following the perturbation. Thus, a simplified bacterial community may be able to recover colonization resistance to patients treated with antibiotics.

## Introduction

The process by which gut bacteria prevent *Clostridioides difficile* and other pathogens from infecting and persisting in the intestine is known as colonization resistance (1). Antibiotic-induced disruption of the gut bacterial community breaks down colonization resistance and is a major risk factor for *C. difficile* infection (CDI) (2). Gut bacteria inhibit *C. difficile* through the production of bacteriocins, modulation of available bile acids, competition for nutrients, production of short-chain fatty acids, and altering the integrity of the mucus layer (1). After the initial CDI is cleared via antibiotics, patients can become reinfected. When CDI recurs more than once, the gut bacterial community from a healthy person typically is used to restore the gut community in the patient with recurrent CDI (3). Fecal microbiota transplant (FMT) is effective, but 10 - 20% of people that receive a FMT will still have another CDI (4). Additionally, transfer of a whole fecal community can incidentally transfer pathogens and cause adverse outcomes (5). While the FMT is effective at curing recurrent CDI, it also has risks that must be considered.

The benefits and risks of the FMT has led to the development of simplified bacterial communities to treat CDI. Synthetic communities are more defined than an FMT, making them easier to regulate as a drug. Tvede and Rask-Madsen were the first to successfully treat CDI with a community of isolates cultured from human feces (6). More recently, Lawley *et al*. analyzed murine experiments and the fecal communities from patients with CDI to develop a synthetic community of six isolates to inhibit *C. difficile* colonization (7). Simplified communities derived from human fecal communities, by methods such as selective isolation of spores or culturing bacteria, have cured recurrent CDI in their initial application (8–10). Although, a recent phase 2 trial of SER-109, a spore-based treatment, failed its phase 2 clinical trial (11), these therapies have the potential to offer the benefits of the FMT without the associated risks but are only used once a patient has had multiple CDIs. For these to be successful, we need a better approach to identify candidate bacterial populations. Recently, an autologous FMT was shown to be effective at restoring the gut microbiota in allogeneic hematopoietic stem cell transplantation patients and prevented future complications, such as systemic infections (12). It is unclear whether an treatment similar to an autologous FMT or simplified bacterial communities could be used to restore susceptible communities and prevent CDI (13).

Because FMT is often sufficient to restore colonization resistance to people with a current infection, we hypothesized that a fecal community should be sufficient to restore colonization resistance to an uninfected community. Therefore, we tested whether a fecal community transplant (FCT) pre-treatment would prevent or clear *C. difficile* colonization and how variation in susceptibility to *C. difficile* infection would affect the effectiveness of FCT pre-treatment. After testing the same FCT pre-treatment across different antibiotic-induced susceptibilities, we sought to determine whether diluted FCT pre-treatment could maintain the inhibition of *C. difficile* colonization and identify the bacterial populations associated with colonization resistance and clearance.

## Results

### Effect of fecal transplant on *C. difficile* colonization was not consistent across antibiotic treatments

Our previous research demonstrated that when mice were perturbed with different antibiotics, there were antibiotic-specific changes to the microbial community that resulted in different levels of colonization and clearance of *Clostridioides difficile* infection (14). Because each of these treatments opened different niche spaces that *C. difficile* could fill, we hypothesized that the resulting community varied in the types of bacteria required to recover colonization resistance. To test the ability of the murine communities to recover colonization resistance, we treated conventionally raised SPF C57BL/6 mice with either clindamycin, cefoperazone, or streptomycin. After a short recovery period, the mice were given either phosphate-saline buffer (PBS) or a fecal community transplant via oral gavage (Figure 1). The fecal community was obtained from untreated mice. One day after receiving the FCT, the mice were challenged with 10^3^ *C. difficile* 630 spores. One day after the challenge, mice that were treated with either clindamycin or cefoperazone and received the FCT pre-treatment had similar amounts of *C. difficile* colony forming units (CFU) as those which received PBS. Among the clindamycin-treated mice, *C. difficile* colonization was cleared at similar rates regardless of whether they received the FCT or PBS pre-treatments (Figure 2). For cefoperazone-treated mice, *C. difficile* colonized all of the mice, but the mice that received the FCT pre-treatment cleared the infection (Figure 2). For the streptomycin-treated mice, the FCT pre-treatment resulted in either no detectable *C. difficile* colonization (8 of 14) or an infection that the community cleared within 5 days (Figure 2). For mice that would have normally had a persistent infection, the FCT enabled them to clear the infection and in the streptomycin-treated mice it was able to prevent infection entirely for some mice.

**Figure 1.**
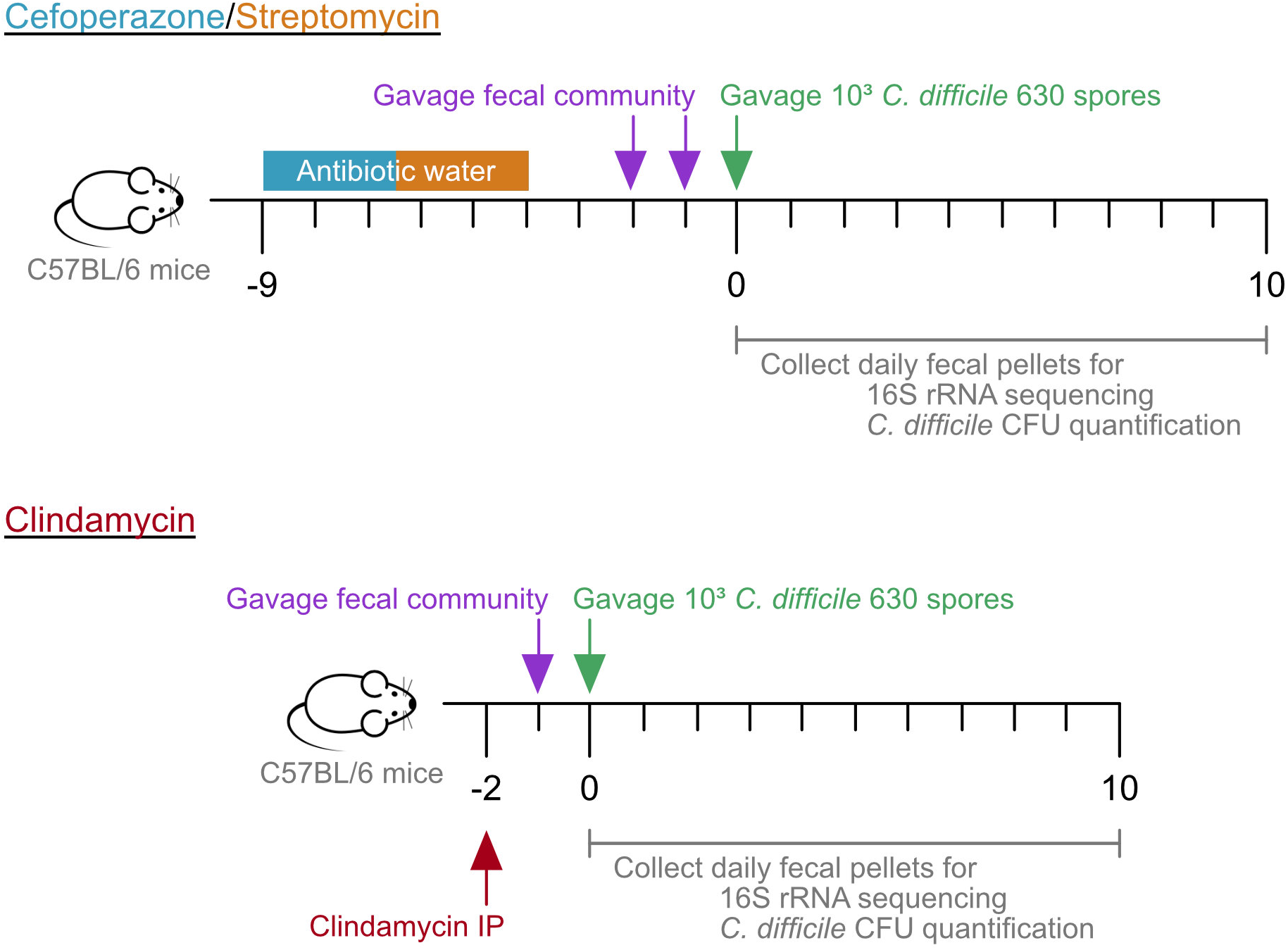
Mouse experiment timeline. Mice were given water with cefoperazone (0.5 mg/ml) or streptomycin (5 mg/ml) for 5 days. The mice were returned to untreated water for the remainder of the experiment. Two days after the antibiotic water was removed, mice were orally gavaged 100 *μ*l of PBS or fecal community, once a day for two days. The following day, the mice were challenged with 10^3^ *C. difficile* 630 spores. Alternatively, mice were given an intraperitoneal injection of clindamycin (10 mg/kg) 2 days prior to *C. difficile* infection. 24 hours later, mice were orally gavaged with 100 *μ*l of PBS or fecal community. The following day, the mice were challenged with 10^3^ *C. difficile* 630 spores. Fecal pellets were collected prior to treatment (day −9 for cefoperazone/streptomycin, day −2 for clindamycin), cessation of antibiotics (day −2 for cefoperazone/streptomycin, day −1 for clindamycin), prior to *C. difficile* infection (day 0), and each of the following 10 days.

**Figure 2.**
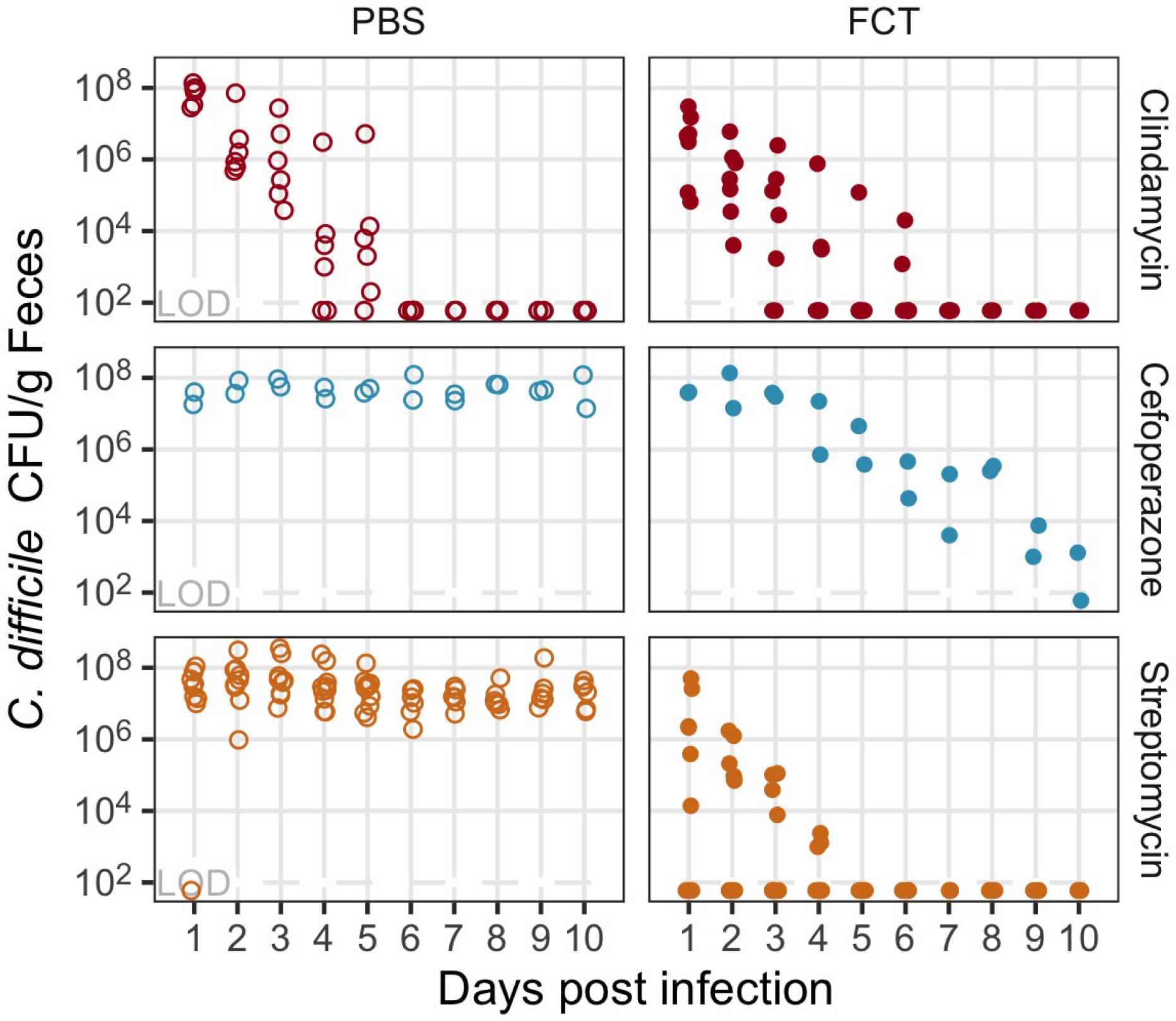
Fecal community transplant inhibited *C. difficile* colonization for mice treated with cefoperazone or streptomycin. *C. difficile* CFU per gram of feces for mice treated with clindamycin (red points), cefoperazone (blue points), or streptomycin (orange points). Mice were orally gavaged either PBS (open circles) or FCT (fecal community transplant, filled circles) prior to the *C. difficile* infection. Each point represents an individual mouse. LOD = limit of detection.

### Diluted fecal communities prevented colonization and promoted clearance for streptomycin-treated mice

Next, we sought to test whether mice that received a diluted FCT pre-treatment could still benefit. To identify the minimally effective dilution, we repeated the same experimental design (Figure 1) with the FCT diluted serially down to 1:10^5^. Since the FCT pre-treatment had no detected effect in clindamycin-treated mice, we did not study those mice further. Cefoperazone-treated mice pre-treated with diluted FCT, 1:10 and lower, were not affected and were colonized throughout the experiment (Figure 3). Streptomycin-treated mice pre-treated with diluted FCT either regained colonization resistance or were enabled to clear *C. difficile*. The streptomycin-treated mice pre-treated with FCT as dilute as 1:10^3^ cleared *C. difficile*. The streptomycin-treated mice pre-treated with FCT as dilute as 1:10^2^ had no *C. difficile* CFU detected throughout the length of the experiment. While more mice pre-treated with lower FCT dilutions were colonized (FCT 6 of 14 were colonized, 1:10 10 of 12 were colonized, 1:10^2^ 10 of 14 were colonized), the colonized mice that received the lower dilutions were still able to clear *C. difficile* (Figure S1). Thus, the simplified fecal communities from the diluted FCT were able to restore colonization resistance and promote clearance of *C. difficile* in streptomycin-treated mice.

**Figure 3.**
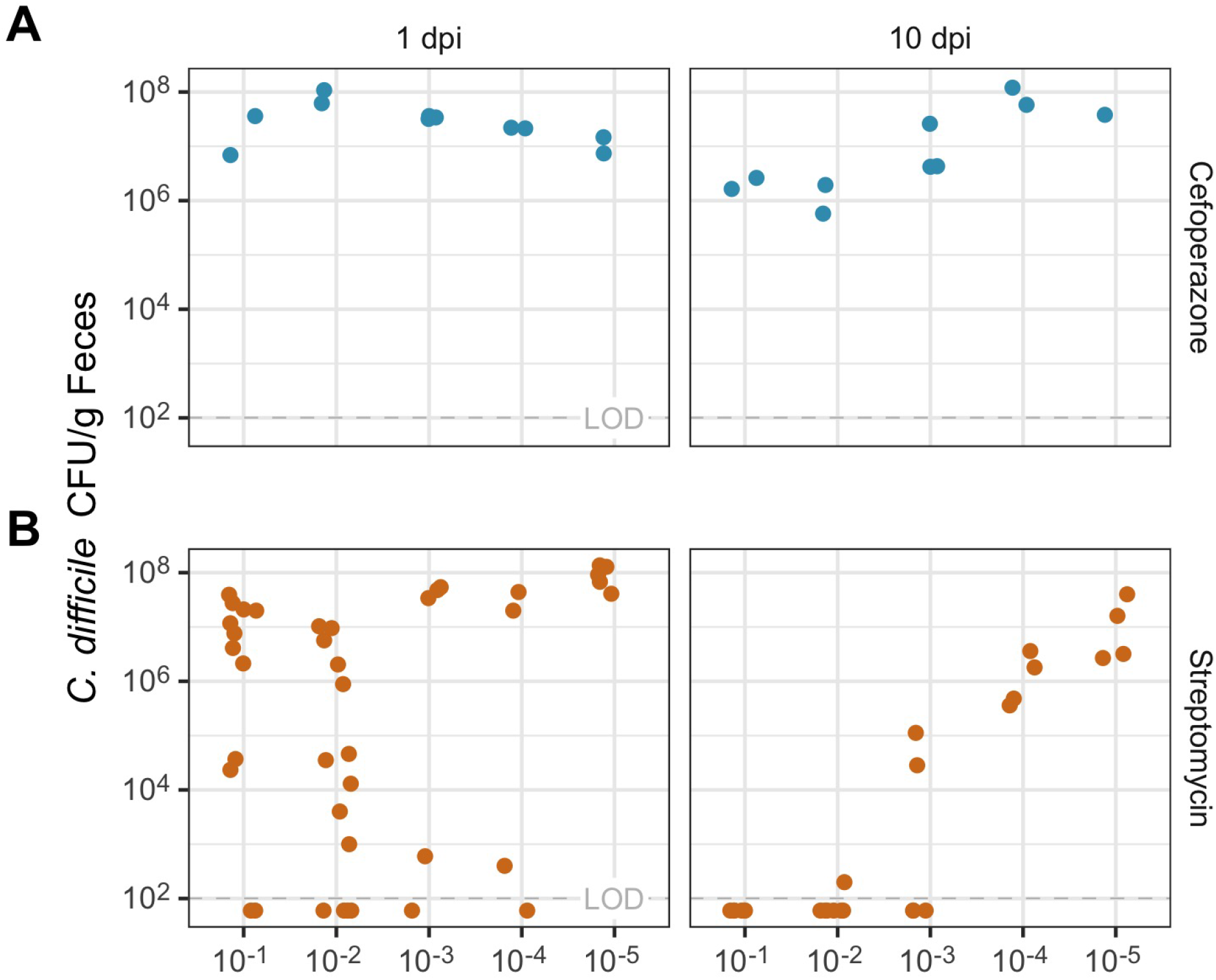
Diluted FCT inhibited *C. difficile* colonization for mice treated with streptomycin. *C. difficile* CFU per gram of feces for mice treated with cefoperazone (blue points) or streptomycin (orange points). Mice were orally gavaged with a dilution of FCT (1:10 to 1:10^5^) prior to the *C. difficile* infection at (A) one day post *C. difficile* infection (dpi) and (B) 10 dpi. Each point represents an individual mouse. LOD = limit of detection.

The simplified fecal communities of the diluted FCT may have reduced abundance and membership. We compared the FCT communities to determine the differences between the dilutions. The most significant difference between the communities of the FCT and its dilutions was the quantity of the 16S rRNA gene, which decreased monotonically (Figure S2D). The FCT dilutions of 1:10^3^ to 1:10^5^) of the had few samples with sufficient sequencing depth to provide bacterial community information. The FCT and its dilutions were not significantly different in either *α*-diversity (number of operational taxonomic units (OTUs) (S_obs_) or Inverse Simpson) or *β*-diversity (*θ*_YC_) (Figure S2A-C). Populations of *Acetatifactor, Enterobacteriaceae, Lactobacillus, Ruminococcaceae*, and *Turicibacter* correlated with the FCT dilution factor (Figure S2E). Overall, the abundance of the bacteria appeared to be largest difference between FCT and its dilutions.

### Murine gut bacterial communities had not recovered their diversity by the time of *C. difficile* challenge

To elucidate the effects of the fecal community dilution on the murine gut bacterial community and *C. difficile* infection, we sequenced the V4 region of the 16S rRNA gene from the fecal community. For the gut communities, in comparison to the initial community (day −9), FCT pre-treatment did not result in a significant recovery of diversity at the time of *C. difficile* challenge (day 0) for cefoperazone-treated mice (Figure S3) or streptomycin-treated mice (Figure 4). At the end of the experiment (day 10), the gut bacterial communities were more similar to their initial community in *α*-diversity (number of OTUs (S_obs_) and Inverse Simpson diversity index) and *β*-diversity (*θ*_YC_) diversity. The mice pre-treated with less dilute FCT were most similar to the initial community structure, whereas, the mice pre-treated with more dilute FCT resulted in little recovery of diversity, similar to the mice given PBS. Thus, the effect of FCT was not large enough at the time of *C. difficile* challenge to be detected in the community diversity but sufficient to affect *C. difficile* colonization. This would suggest the effect was driven by the most abundant populations.

**Figure 4.**
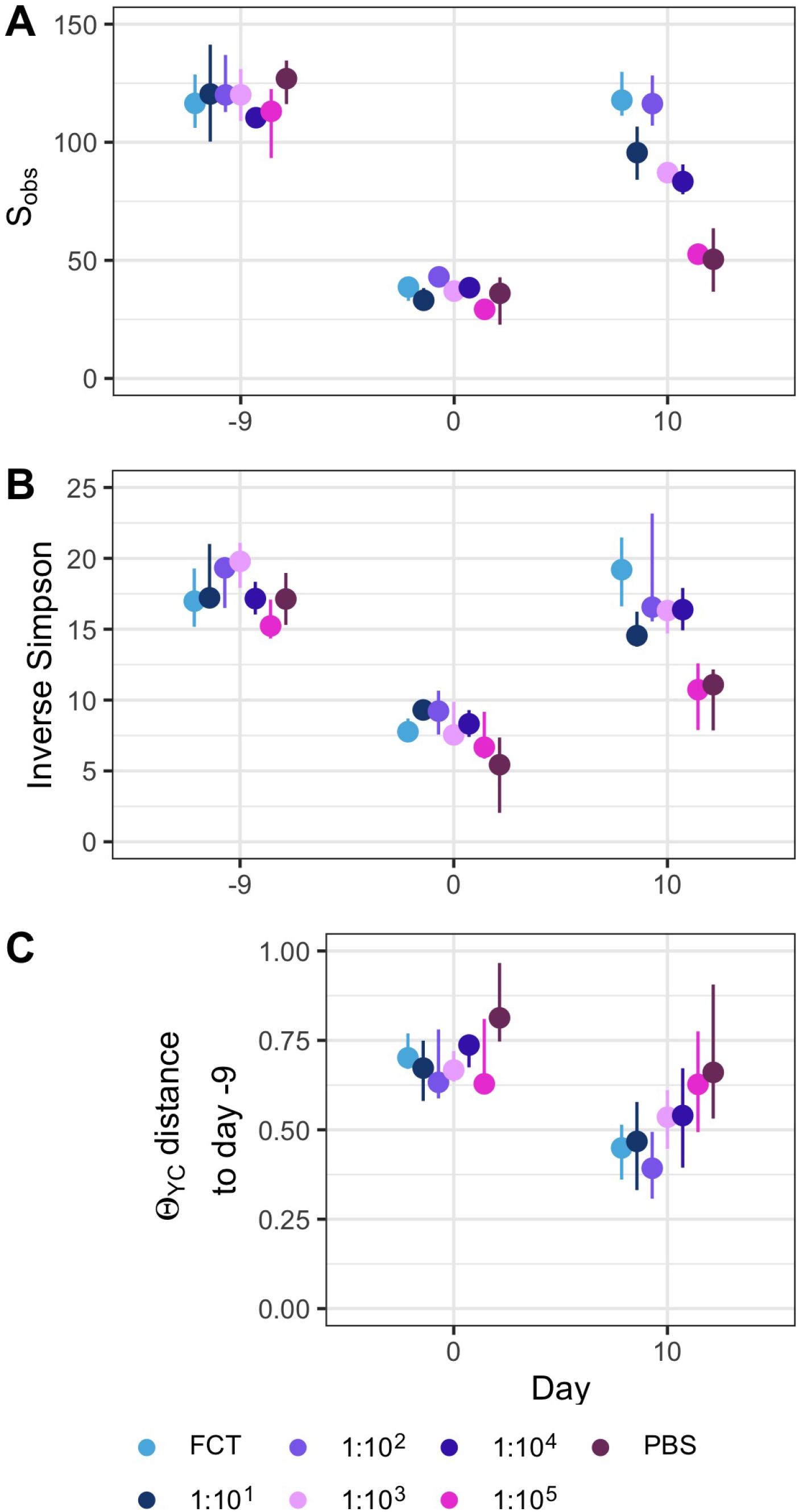
Diversity of murine gut bacterial community had not recovered at the time of *C. difficile* infection in streptomycin-treated mice. *α*-diversity, measured by S_obs_ (A) and Inverse Simpson (B), prior to beginning antibiotic treatment (day −9), after fecal community transplant on the day of *C. difficile* challenge (day 0) and at the end of the experiment (day 10). (C) *β*-diversity, measured by *θ*_YC_, distance between community structures on day 0 or 10 compared to the community prior to antibiotic treatment (day −9) community of that individual. Data are grouped by the transplant received, undiluted fecal community (FCT), diluted fecal community (1:10^1^-1:10^5^), or PBS. Points are median values and lines represent the interquartile range.

### Gut bacterial community members are differentially abundant in streptomycin-treated mice resistant to colonization

Although there were no significant differences in diversity at the time of challenge, we next investigated how the individual bacterial populations were different in the uncolonized streptomycin-treated mice pre-treated with FCT. We used linear discriminant analysis (LDA) effect size (LEfSe) analysis to identify OTUs within the fecal bacterial communities from the streptomycin-treated mice that were differentially abundant between uncolonized and colonized mice. The antibiotic treatment significantly altered 99 OTUs (Figure S5), but on the day of *C. difficile* challenge only 7 OTUs were differentially abundant between colonized and uncolonized communities (Figure 5A). Communities resistant to *C. difficile* colonization had more abundant populations of OTUs related to *Akkermansia, Clostridiales, Olsenella*, and *Porphyromonadaceae* and less abundant populations of an OTU related to *Enterobacteriaceae*. Thus, a small portion of OTUs, relative to the changes due to streptomycin treatment, were differentially abundant in mice that resisted *C. difficile* colonization compared to those that were colonized.

**Figure 5.**
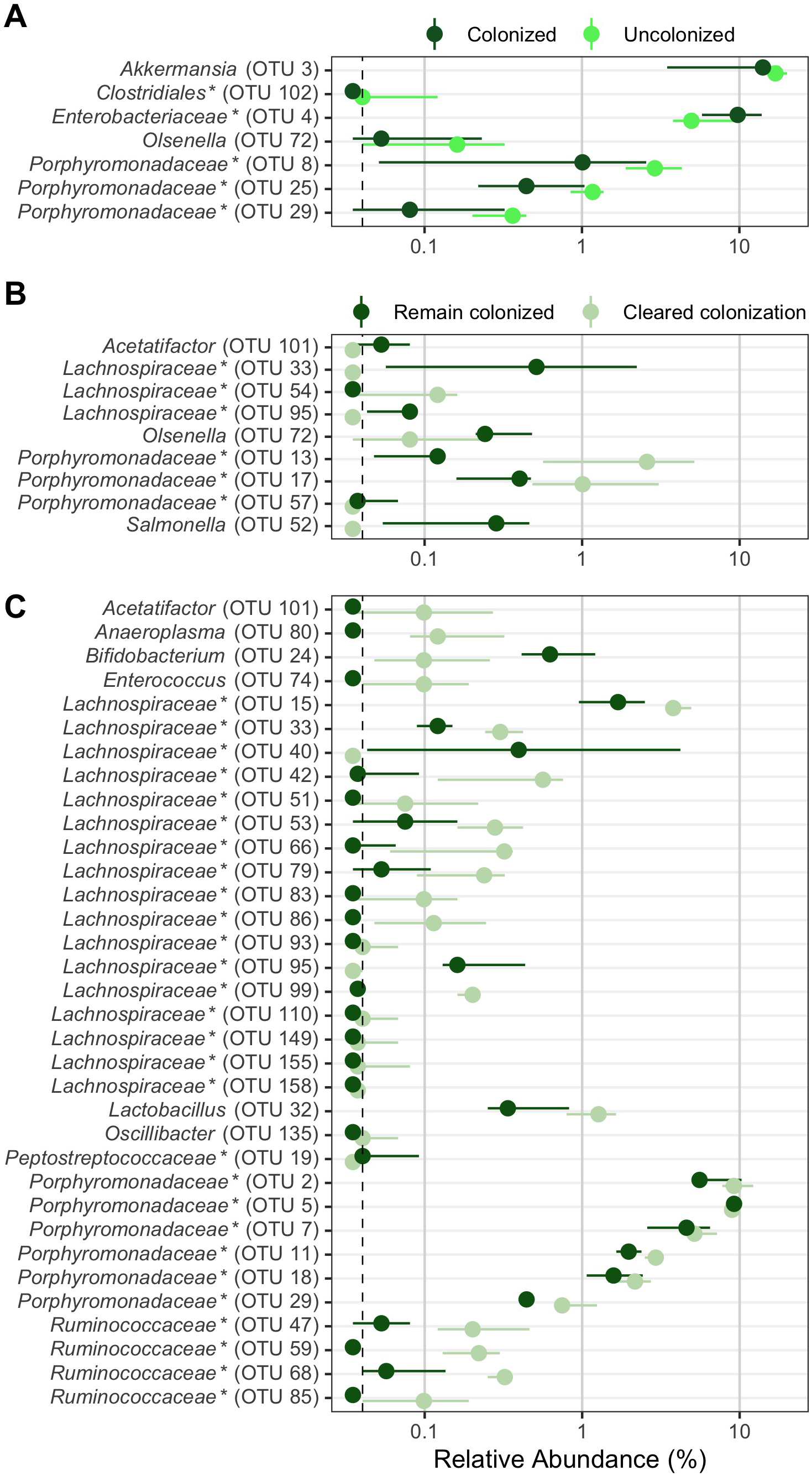
Bacterial community OTUs differentially abundant in streptomycin-treated mice which resisted or cleared colonization. Murine gut bacterial community OTUs that were significantly different by LEfSe analysis. OTUs from streptomycin-treated mice at the time of *C. difficile* challenge (day 0) which were differentially abundant between (A) mice that were colonized (dark green) and those that were not (no detectable CFU throughout the experiment, bright green) or (B) mice that remained colonized (dark green) and those that cleared colonization (CFU reduced to below the limit of detection by the end of the experiment, faint green). (C) OTUs from streptomycin-treated mice at the end of the experiment (day 10) which were differentially abundant between mice that remained colonized (dark green) and those that cleared colonization (CFU reduced to below the limit of detection by the end of the experiment, faint green). Points are median values and lines represent the interquartile range. Dashed vertical line is the limit of detection. OTUs ordered alphabetically. * indicates that the OTU was unclassified at lower classification rank.

### Murine gut bacterial communities that cleared *C. difficile* colonization were more similar to the initial community

To better understand the differences in streptomycin-treated murine fecal community that contributed to *C. difficile* clearance, we compared the communities that cleared *C. difficile* to those that did not at the time of challenge and 10 days post infection. Communities from mice that cleared colonization were more similar to their initial community at the end of the experiment than the mice that remained colonized (Figure S4). At the time of *C. difficile* challenge, 9 OTUs were differentially abundant between communities that remained colonized to those that cleared colonization (Figure 5B). Communities that cleared *C. difficile* colonization had more abundant populations of OTUs related to *Porphyromonadaceae* and *Lachnospiraceae* and less abundant populations of OTUs related to *Acetatifactor, Lachnospiraceae, Olsenella, Porphyromonadaceae*, and *Salmonella*. At the end of the experiment, 29 of the 34 differentially abundant OTUs were more abundant in the mice that were able to clear the colonization (Figure 5C). The relative abundance of OTUs related to *Acetatifactor, Anaeroplasma*, *Enterococcus*, *Lachnospiraceae*, *Lactobacillus*, *Porphyromonadaceae*, and *Ruminococcaceae* were higher in communities that cleared, recovering in abundance from the streptomycin treatment. Multiple OTUs related to *Lachnospiraceae* and *Porphyromonadaceae* (N = 14 and N = 5, respectively) were significant and accounted for greater portions of the community (more than 10%). However one *Porphyromonadaceae* population (OTU 5) and two *Lachnospiraceae* related populations (OTU 40 and 95) were more abundant in the mice that remain colonized. Thus, the more the gut bacterial members returned to their initial abundance, there was a greater likelihood of clearing *C. difficile*.

### Negative associations dominated the interactions between the gut bacterial community and *C. difficile* in streptomycin-treated mice

In streptomycin-treated mice, pre-treatment with FCT and its dilutions had different effects on the bacterial community members which resulted in different community relative abundance and *C. difficile* colonization dynamics. We quantified the relationships occurring throughout this experiment using SPIEC-EASI (sparse inverse covariance estimation for ecological association inference) to construct a conditional independence network. Here, we focused on the associations of the *C. difficile* subnetwork (Figure 6). *C. difficile* CFU had positive associations with populations of OTUs related to *Enterobacteriaceae* (OTU 4) and *Peptostreptococcaceae* (OTU 19). OTUs related to *Clostridiales* (OTU 27), *Lachnospiraceae* (OTUs 15, 51, and 83), and *Porphyromonadaceae* (OTUs 23, 25, and 29) had negative associations with *C. difficile*, as well as the OTUs related to *Enterobacteriaceae*, and *Peptostreptococcaceae*. Overall, the majority of the associations between *C. difficile* and the gut bacterial community in streptomycin-treated mice were negative. This suggests this subset of the community may be driving the inhibition of *C. difficile* in streptomycin-treated communities.

**Figure 6.**
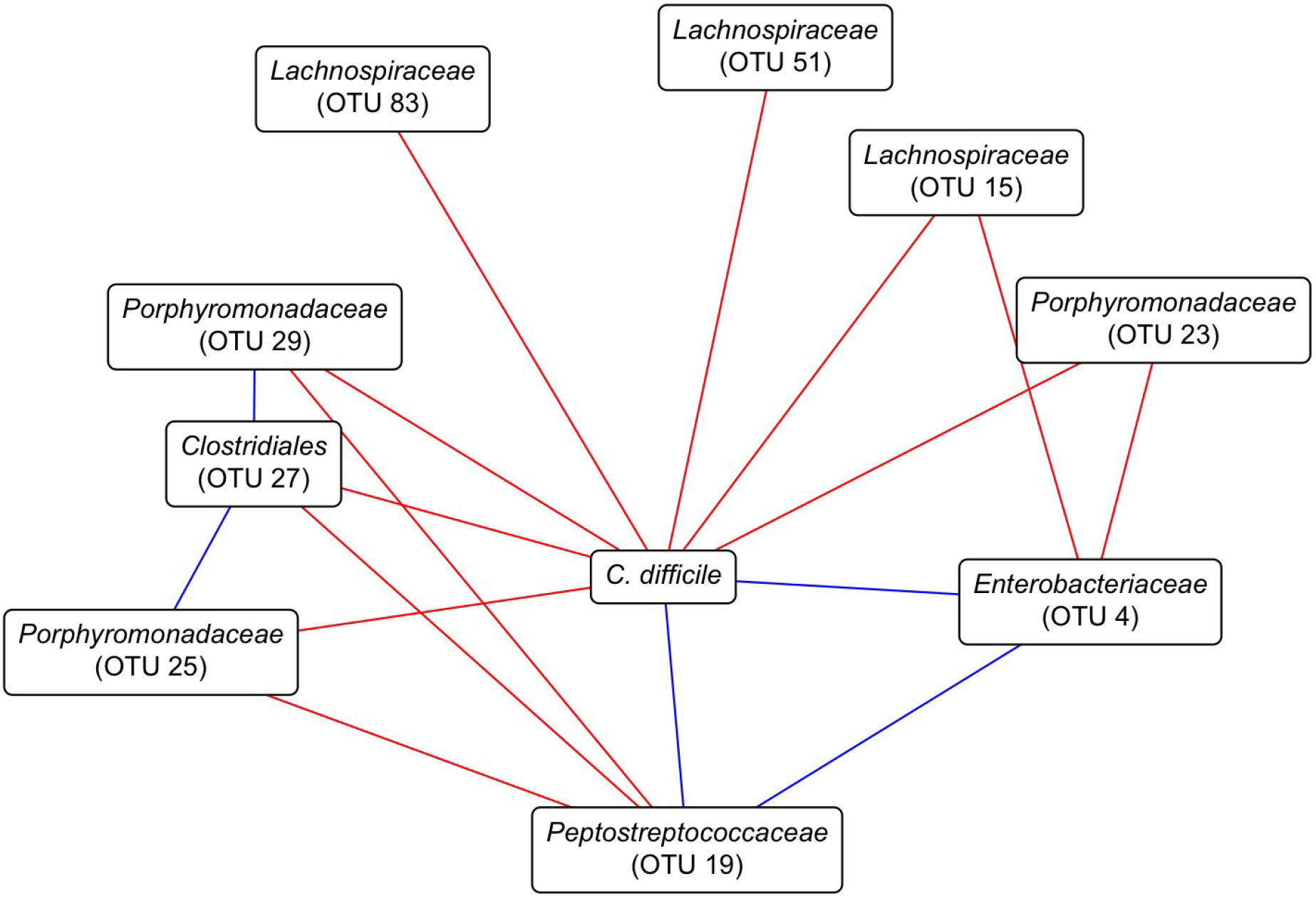
Streptomycin-treated murine fecal community associations with *C. difficile*. Network constructed with SpiecEasi from the OTU relative abundances and *C. difficile* CFU data from 1 through 5 days post *C. difficile* infection. Red lines represent negative associations and blue lines indicate positive associations. *C. difficile* is based on CFU counts and *Peptostreptococcaceae* (OTU 19), the OTU most closely related to *C. difficile*, is based on sequence counts. Only *C. difficile* subnetwork shown.

## Discussion

Transplanting the fecal community from untreated mice to antibiotic-treated mice prior to being challenged with *C. difficile* varied in effectiveness based on the antibiotic treatment. This indicated that FCT pre-treatment can prevent *C. difficile* colonization in an antibiotic-specific manner. Additionally, by diluting the FCT we were able to narrow the community changes responsible for the effect to the most abundant OTUs. Overall, these results show that a simplified fecal community can assist a perturbed microbiota in preventing or resisting *C. difficile* colonization but the effect was dependent on the antibiotic that was given.

By diluting the FCT we were able to narrow the definition of the minimal community features that restored colonization resistance. Bacterial interactions with *C. difficile* were associated with the identity, abundance, and functions of adjacent bacteria. Ghimire *et al*. recently showed individual species that inhibited *C. difficile* in co-culture but when other inhibitory species were added the overall effect on *C. difficile* was changed, in some cases to a increased *C. difficile* growth (15). Based on these observations from their bottom-up approach, it is unclear how more complex combinations would affect the inhibition of *C. difficile*. So instead, we sought to find an inhibitory community using a top-down approach and begin with an inhibitory community. In a recent top-down approach, Auchtung *et al*. recently developed a set of simplified communities from human fecal communities that were grown in minibioreactor arrays and tested for inhibition first *in vitro* then in a mouse model (16). They found four simplified communities that were able to reduce *C. difficile* colonization but with varied effect in a mouse model with the same gut microbiota. One way they simplified the community was through diluting the initial fecal sample. In our experiments, we began with a fresh whole fecal community to first determine if inhibition was possible. In the conditions which *C. difficile* was inhibited, with cefoperazone and streptomycin, we diluted the FCT to determine the minimal community which maintained inhibition. Cefoperazone-treated mice were unable to maintain inhibition of *C. difficile* with diluted FCT pre-treatments and *C. difficile* remained colonized. Streptomycin-treated mice were able to maintain inhibition with diluted FCT pre-treatment. While the diluted FCTs had similar diversity and bacterial abundances, the differences in effect on *C. difficile* revealed the minimal changes associated with either colonization resistance or clearance.

We previously hypothesized that mice treated with either clindamycin, cefoperazone, or streptomycin would not have the same bacterial community changes associated with *C. difficile* clearance (14). In that set of experiments, the dose of the antibiotic was varied to titrate changes to the community and determine what changes allow *C. difficile* to colonize and then be spontaneously cleared. We observed antibiotic-specific changes associated with *C. difficile* clearance. The data presented here complement those observations. For clindamycin-treated mice, there was no difference in colonization, clearance or relative abundance between PBS and FCT pre-treatment. *C. difficle* had similar colonization dynamics. It is possible that there was insufficient time for the FCT to engraft. However, when we added more time between clindamycin treatment and *C. difficile* challenge, *C. difficile* was unable colonize (data not shown). Therefore, clindamycin-treated mice appeared to have been naturally recovering inhibition to *C. difficile*, which was unaffected by the FCT pre-treatment. For cefoperazone-treated mice, the FCT pre-treatment enabled the gut microbiota to eliminate the colonization but only in its most concentrated dose. This observation supports our previous report, indicating that the cefoperazone-treated community is more sensitivity to the amount of FCT it receives since cefoperazone reduced many bacterial groups and associations (Figure S6). As we previously hypothesized, streptomycin-treated mice were enabled to clear with a subset of the community, with the FCT pre-treatment diluted 1:10^3^. Since we titrated the FCT dilutions, we could compare the bacterial communities of the mice which gained the ability to clear *C. difficile* to the mice that received the next dilution which could not to elucidate the minimal relative abundance differences. In agreement with previous studies, OTUs related to *Lachnospiraceae, Porphyromonadaceae*, and *Ruminococcaceae* increased with the clearance of *C. difficle* in the streptomycin-treated mice (14, 17–21). These data agree with our previous hypothesis that a simplified fecal community would only be able to promote clearance of *C. difficile* in streptomycin-treated mice.

In addition to clearing *C. difficile*, a simplified fecal community restored colonization resistance to streptomycin-treated mice. Mice that received FCT pre-treatment as dilute as 1:10^2^ were not colonized to a detectable level. While restoring colonization resistance is not novel (22), here we have shown that the restoration of colonization resistance is dependent on the community perturbation and the fecal community being transplanted. As we identified community members associated with clearance, OTUs related to *Akkermansia*, *Olsenella*, and *Porphyromonadaceae* were more abundant and an OTU related to *Enterobacteriaceae* was less abundant at the time of *C. difficile* challenge. *Enterobacteriaceae* has been associated with *C. difficile* colonization and inflammation (14, 23, 24). Larger populations of *Akkermansia* were associated with preventing colonization, which we had previously observed, potentially indicating the maintenance of a protective mucus layer (14, 25–27). Increased populations of a select set of OTUs related to *Porphyromonadaceae* were also more abundant in mice that were resistant to colonization. *Porphyromonadaceae* may be inhibiting *C. difficile* via buytrate and acetate production, which has been associated with successful FMT treatments (28–30). Different populations of OTUs associated with *Porphyromonadaceae* were associated with colonization resistance than with colonization clearance. These colonization resistance-associated OTUs (OTUs 8, 25 and 29) may have OTU-specific functions or dependent abundances of memebrs of the community, such as *Akkermansia* and *Enterobacteriaceae*. With our top-down approach, we reduced the number of gut bacterial community members that were associated with colonization resistance in streptomycin-treated mice.

We have demonstrated that a simplified bacterial community can restore colonization resistance but the effect of the community and the bacteria that colonized was dependent on the specific changes to the community that were caused by each antibiotic. When transplanting the fecal community into antibiotic-induced susceptible mice, only mice treated with streptomycin were able to restore colonization resistance. Previous studies that have identified simplified communities in a murine model using a homogeneous gut microbiota with a bottom-up approach (7, 19). Treatments supplementing the gut microbiota would benefit from being tested in different communities susceptible to CDI. Further research is necessary to characterize the specific niche spaces *C. difficile* of susceptibilities communities and the specific requirements fill those spaces. Then it may be possible to identify people with gut microbiota that are susceptible to CDI and develop targeted simplified bacterial communities to recover colonization resistance and reduce the risk of CDI.

## Materials and Methods

### Animal care

Mice used in experiments were 6- to 13-week old conventionally reared SPF male C57BL/6 mice obtained from a single breeding colony at the University of Michigan. During the experiment, mice we housed with two or three mice per cage. All murine experiments were approved by the University of Michigan Animal Care and Use Committee (IACUC) under protocol number PRO00006983.

### Antibiotic administration

Mice were given either cefoperazone, clindamycin, or streptomycin. Cefoperazone (0.5 mg/ml) and streptomycin (5 mg/ml) were administered via drinking water for 5 days, beginning 9 days prior to *C. difficile* challenge. Antibiotic water was replaced every two days. Clindamycin (10 mg/kg) was injected into the intraperitoneal space, 2 days prior to challenge with *C. difficile*. All antibiotics were filter sterilized with a 0.22 *μ*m syringe filter prior to use.

### Fecal community transplants

Fecal pellets were collected from similar aged C57BL/6 mice not being used in an experiment the day of the fecal community transplants. 15-20 pellets were collected and weighed. The fecal pellets were homogenized weight per weight in phosphate-saline buffer (PBS) containing 15% glycerol (fecal community transplant, FCT) in anaerobic conditions. The FCT was serially diluted in PBS containing 15% glycerol down to 1:10^5^ fecal dilution and aliquoted into tubes for gavaging into mice. One set of aliquots were frozen at −80°C to be used the following day for the cefoperazone and streptomycin experiments. Frozen aliquots were thawed at 37°C for 5 minutes prior to being used. All fecal community dilutions were centrifuged at 7500 RPM for 60 seconds. Mice were inoculated with 100 uL of the fecal dilution oral via a 21 gauge gavage needle. Fecal community transplants were administered from most dilute to least, which began with mice receiving PBS and finished with mice receiving FCT. Aliquots were frozen at −80°C after use for sequencing. These experiments were repeated 8 times with a different starting source each time.

### 16S rRNA quantitative real-time PCR

Quantitative analysis of 16S rRNA in fecal community dilutions used for FCT was carried out using quantitative real-time PCR using primers and cycler conditions specified previously (31). Reaction volumes were prepared using 6 uL of SYBRTM Green PCR Master Mix (Applied Biosciences Ref 4344463), 1 uL each forward and reverse primer, and 2 uL sample DNA template. All qPCR reactions were run on a LightCycler^®^ 96 (Roche Ref 05815916001) using instrument-specific plates and seals.

### *C. difficile* challenge

For experiments using streptomycin or cefoperazone, mice were given untreated drinking water for 96 hours before challenging with *C. difficile* strain 630Δerm spores. For experiments using clindamycin, mice were given untreated drinking water for 48 hours the time of the intraperitoneal injection and being challenged with *C. difficile* strain 630Δerm spores. *C. difficile* spores were aliquoted from a single spore stock stored at 4°C. Spore concentration was determined two days prior to the day of challenge (32). Mice were inoculated with 10^3^ *C. difficile* spores via oral gavage. After inoculating the mice, remaining spore solution was serially diluted and plated to confirm the spore concentration.

### Sample collection

Fecal samples were collected prior to administering antibiotics, after antibiotics were removed, prior to *C. difficile* challenge and on each of the 10 days post infection. Approximately 15 mg of each fecal sample was collected and weighed for plating *C. difficile* colony forming units (CFU) and the remaining sample was frozen at −80°C for later sequencing. The weighed fecal samples were anaerobically serially diluted in PBS, plated on TCCFA plates, and incubated at 37°C for 24 hours. The resultant colonies were enumerated to determine the *C. difficile* CFUs (33).

### DNA sequencing

Total bacterial DNA was extracted from the frozen samples by the MOBIO PowerSoil-htp 96-well soil DNA isolation kit. We amplified the 16S rRNA gene V4 region and the amplicons were sequenced on an Illumina MiSeq as described previously (34).

### Sequence curation

Sequences were processed using mothur (v.1.44.1) (34, 35). We used a 3% dissimilarity cutoff to group sequences into operational taxonomic units (OTUs) and a naive Bayesian classifier with the Ribosomal Database Project training set (version 16) to assign taxonomic classifications to OTUs (36). We sequenced a mock community of a known 16S rRNA gene sequences and composition. We processed this mock community in parallel with our samples to determine the error rate for our sequence curation, which resulted in an error rate of 0.029%.

### Statistical analysis and modeling

We calculated diversity metrics in mothur. For *α*-diversity comparisons, we calculated the number of OTUs (S_obs_) and the Inverse Simpson diversity index. For *β*-diversity comparisons, we calculated dissimilarity matrices based on metric of Yue and Clayton (*θ*_YC_) (37). We rarefied samples to 2,480 sequences per sample to limit uneven sampling biases. OTUs were subsampled to 2,480 counts per sample. We tested for differences in relative abundance between outcomes with LEfSe in mothur. All other statistical analyses and data visualization was completed in R (v4.0.5) with the tidyverse package (v1.3.1). Pairwise comparisons of *α*-diversity (S_obs_ and Inverse Simpson), *β*-diversity (*θ*_YC_), were calculated by pairwise Wilcoxon rank sum test. Correlations between bacterial genera and fecal community dilution were calculated using the Spearman correlation. *P* values were corrected for multiple comparisons with a Benjamini and Hochberg adjustment for a type I error rate of 0.05 (38). For streptomycin experiments, conditional independence networks were calculated from the day 1 through 5 samples of all mice using SPIEC-EASI (sparse inverse covariance estimation for ecological association inference) methods from the SpiecEasi R package after optimizing lambda to 0.001 with a network stability between 0.045 and 0.05 (v1.0.7) (39).

## Code availability

Scripts necessary to reproduce our analysis and this paper are available in an online repository (https://github.com/SchlossLab/Lesniak_restoreCR_XXXX_2022).

## Sequence data accession number

All 16S rRNA gene sequence data and associated metadata are available through the Sequence Read Archive via accession SRP373949.

## Acknowledgements

This work was supported by several grants from the National Institutes for Health R01GM099514, U19AI090871, U01AI12455, and P30DK034933. Additionally, NAL was supported by the Molecular Mechanisms of Microbial Pathogenesis training grant (NIH T32 AI007528). The funding agencies had no role in study design, data collection and analysis, decision to publish, or preparation of the manuscript.

## Author contributions

Conceptualization: N.A.L., S.T., P.D.S.; Data curation: N.A.L., L.B., K.M.; Formal analysis: N.A.L.,; Investigation: N.A.L., S.T., A.H., A.T., J.C., L.B., K.M., P.D.S.; Methodology: N.A.L., S.T., P.D.S.; Resources: N.A.L., S.T., L.B., K.M., P.D.S.; Software: N.A.L.; Visualization: N.A.L., P.D.S.; Writing - original draft: NAL; Writing - review & editing: N.A.L., S.T., A.H., A.T., J.C., L.B., K.B., P.D.S.; Funding acquisition: P.D.S.; Project administration: P.D.S.; Supervision: P.D.S.

---

**Figure S1.**
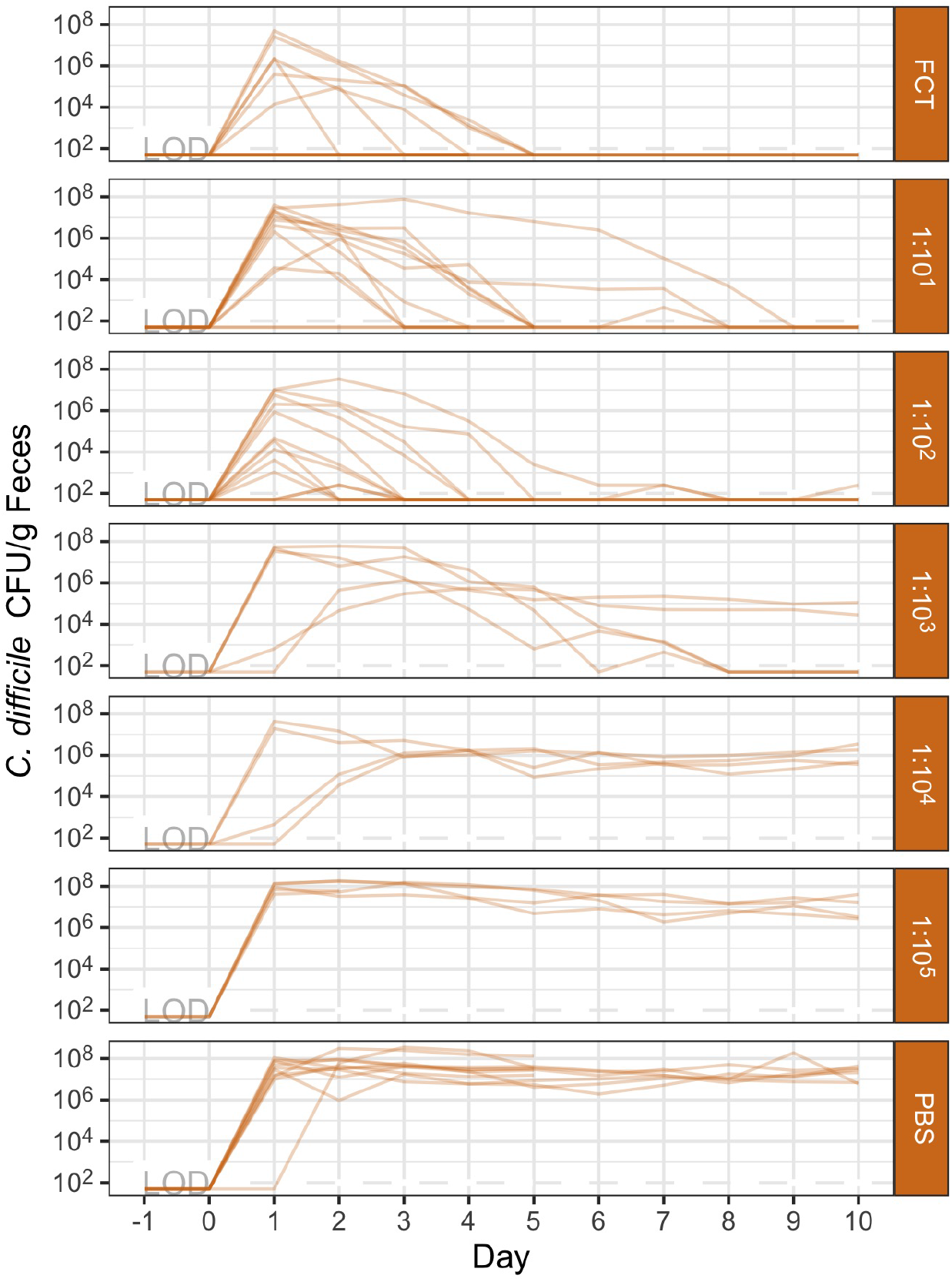
*C. difficile* colonization dynamics in streptomycin-treated mice across all prophylactic transplant treatments. *C. difficile* CFU per gram of feces for streptomycin-treated mice orally gavaged PBS, fecal community transplant (FCT), or diluted FCT (1:10-1:10^5^) prior to the *C. difficile* infection. Each semi-transparent line represents an individual mouse. Mice challenged with 10^3^ *C. difficile* 630 spores on day 0. Lines grouped by the transplant treatment received. LOD = limit of detection.

**Figure S2.**
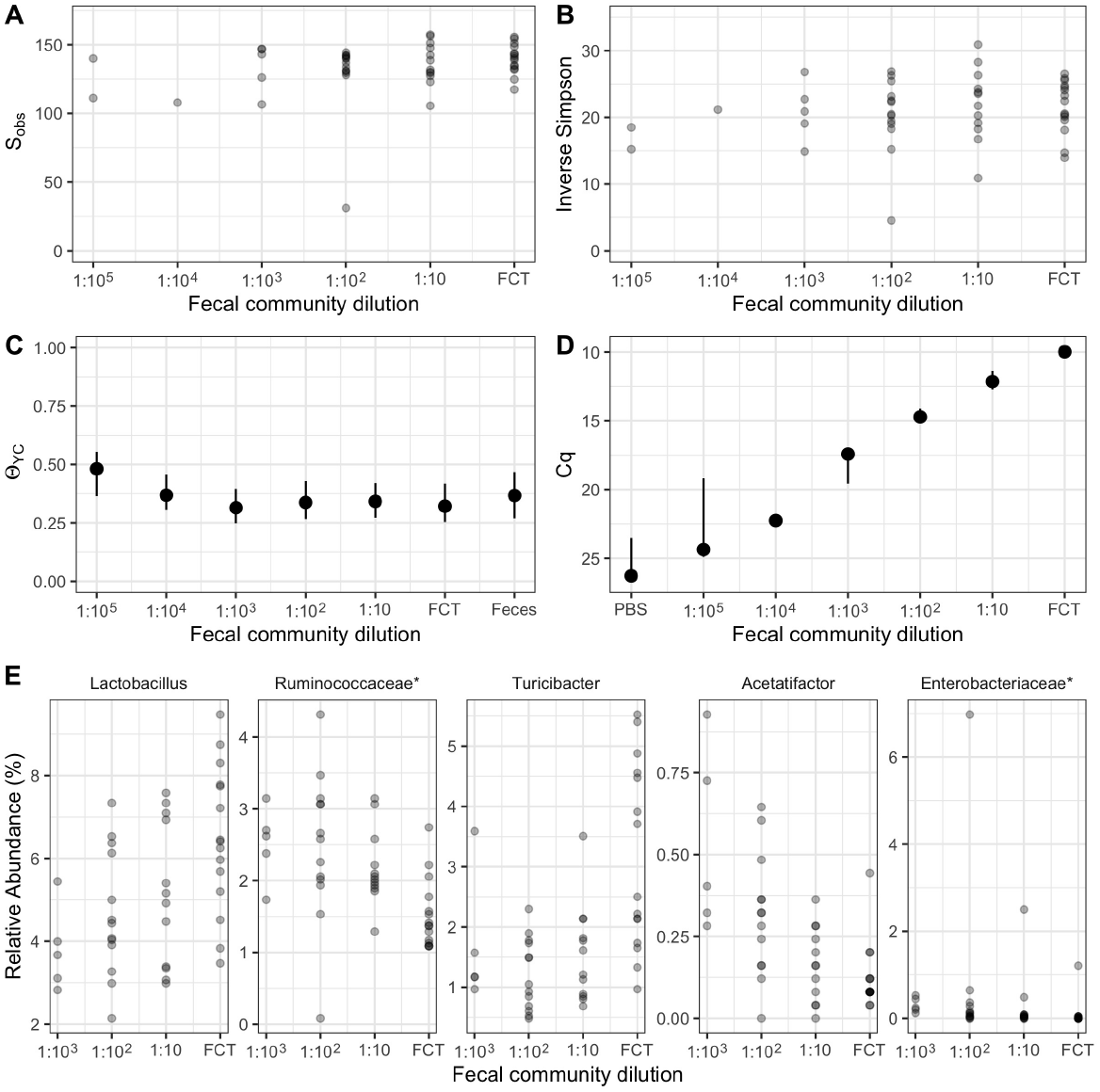
Diversity and quantification of fecal community dilutions used for prophylactic transplants in antibiotic-treated mice. (A-C) Diversity of fecal community dilutions. *α*-diversity, measured by (A) S_obs_ and (B) Inverse Simpson for undiluted fecal community (FCT) and diluted fecal communities (1:10-1:10^5^). Points are individual samples. (C) *β*-diversity, measured by *θ*_YC_, community structure of feces collected from untreated mice, undiluted fecal community (FCT), and diluted fecal communities (1:10-1:10^5^) compared to untreated feces. Points are median values and lines represent the interquartile range. (D) Cq values for qPCR of FCT and its dilutions for eubacterial 16S rRNA gene. Points are median values and lines represent the interquartile range. (E) Relative abundance of bacterial taxonomic groups that significantly correlate with fecal community dilutions (FCT-1:10^3^) by Spearman correlation. Points are individual mice. * indicates that the bacterial taxonomic group was unclassified at lower classification rank.

**Figure S3.**
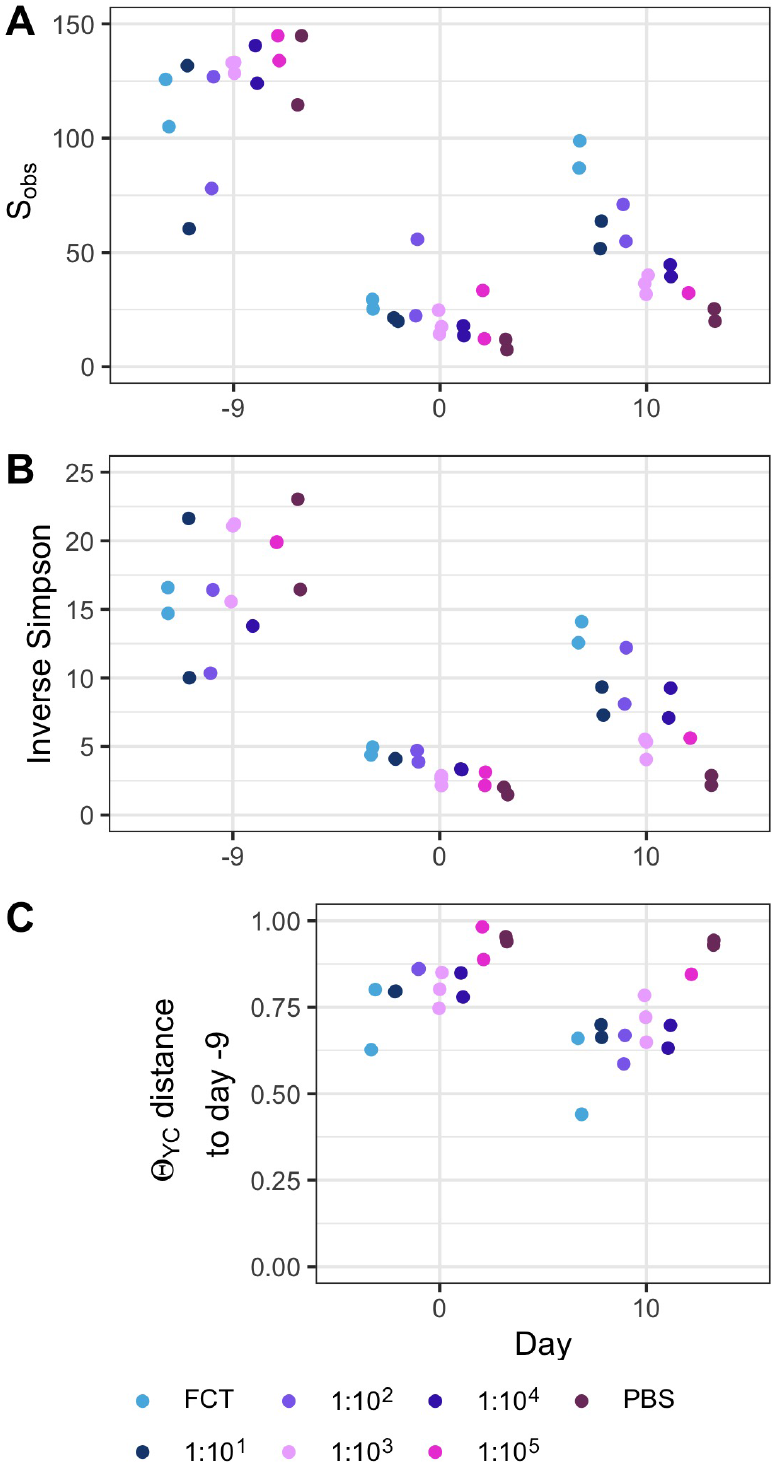
Diversity of murine gut bacterial community was not recovered at the time of *C. difficile* infection in cefoperazone-treated mice. Diversity changes through experiments with cefoperazone-treated mice. *α*-diversity, measured by (A) S_obs_ and (B) Inverse Simpson, prior to beginning antibiotic treatment (day −9), after fecal community transplant on the day of *C. difficile* infection (day 0) and at the end of the experiment (day 10). (C) *β*-diversity, measured by *θ*_YC_, distance between community structures on day 0 or 10 compared to the community prior to antibiotic treatment (day −9) community of that individual. Data are grouped by the transplant received, undiluted fecal community (FCT), diluted fecal community (1:10^1^-1:10^5^), or PBS. Points are individual mice.

**Figure S4.**
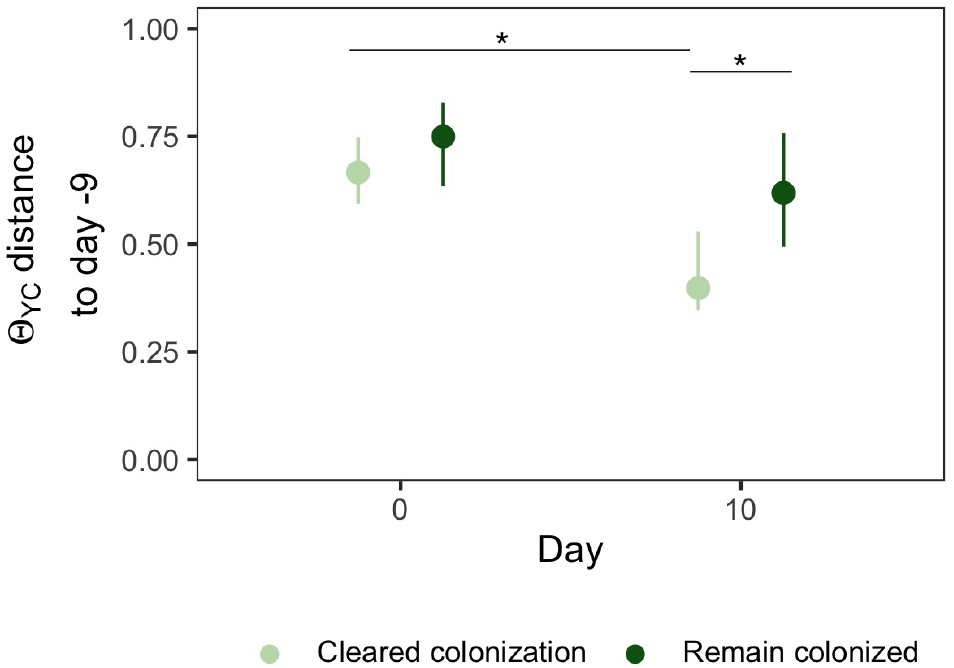
Gut bacterial community of streptomycin-treated mice that cleared colonization were more similar to their initial community. Diversity differences by outcome in streptomycin-treated mice. *β*-diversity, measured by *θ*_YC_, distance between community structures on day 0 or 10 compared to the community prior to antibiotic treatment (day −9) community of that individual. Data are grouped by the outcome, cleared colonization (faint green) or remain colonized (dark green). Points are median values and lines represent the interquartile range. * indicates significant difference by Wilcoxon rank sum test with Bonferroni correction.

**Figure S5.**
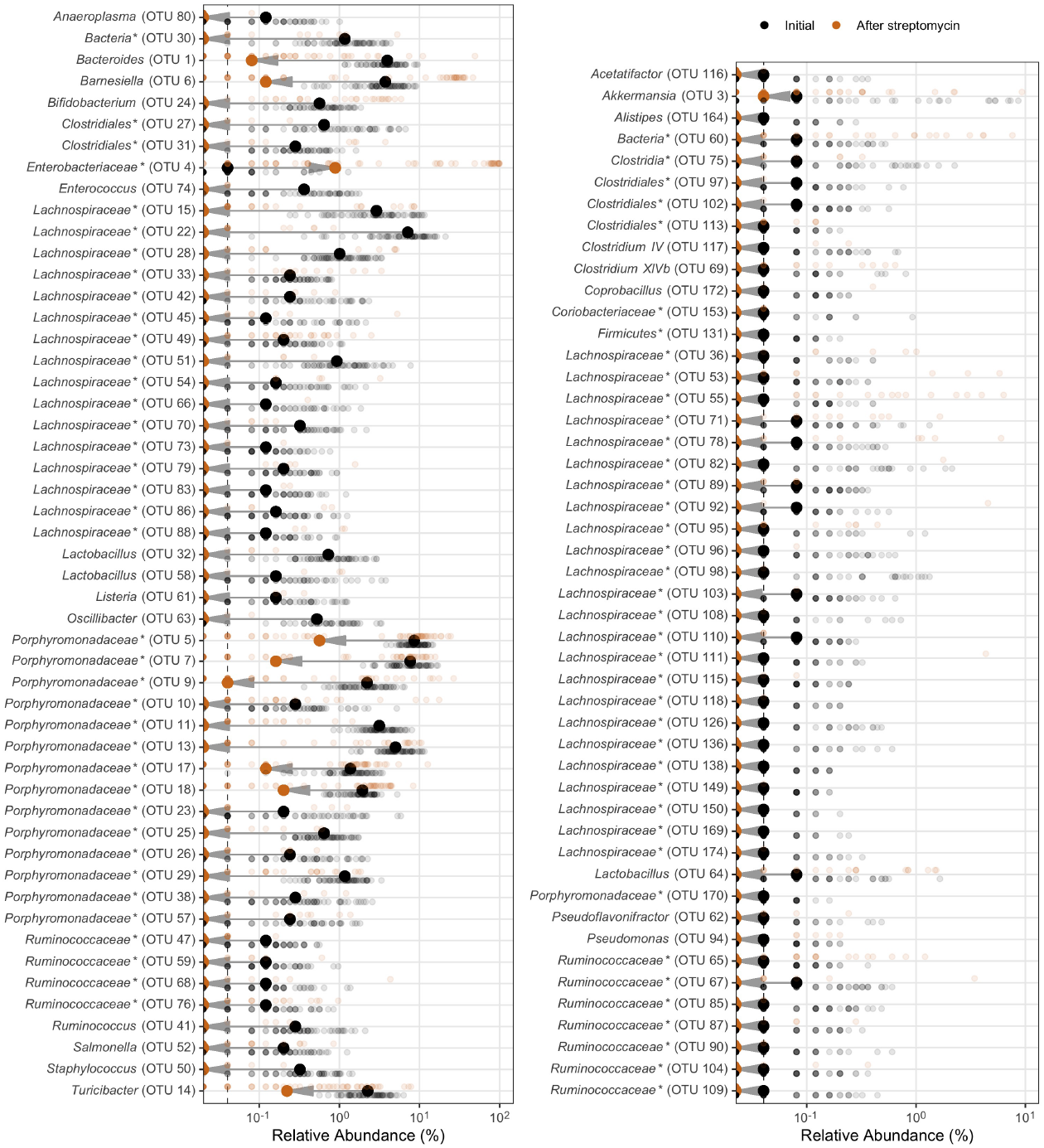
Murine gut bacterial community OTUs differentially abundant with streptomycin treatment. Murine gut bacterial community OTUs that were significantly different by LEfSe analysis between untreated mice (Initial, black) and after 5 days of water with streptomycin (5 mg/ml) and 2 days of untreated water (After streptomycin, orange). Large bold points represent the group median. Small, semi-transparent points represent an individual mouse. Gray arrow indicates the direction the relative abundance shifted with the streptomycin treatment. Left plot displays OTUs with a median relative abundance greater than 0.1%, the OTUs lower are displayed in the right plot. Dashed vertical line is the limit of detection. OTUs ordered alphabetically. * indicates that the OTU was unclassified at lower classification rank.

**Figure S6.**
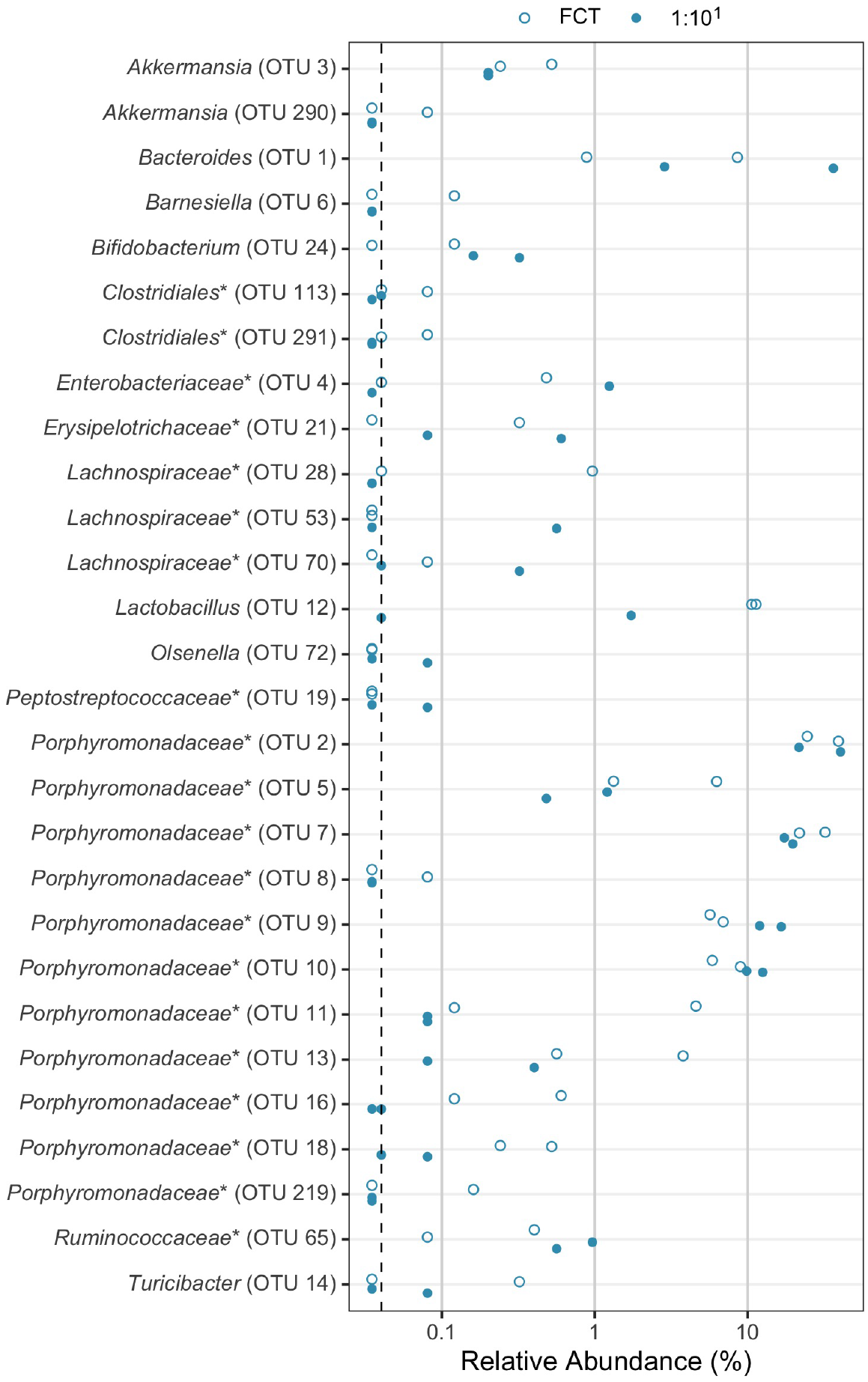
Murine gut bacterial community OTUs of cefoperazone-treated mice at the time of challenge. Murine gut bacterial community OTUs that were present in at least one sample at the time of *C. difficile* challenge (day 0). Mice were pre-treated with either fecal community transplant (FCT, open circles) or FCT diluted 1:10 (filled circles). Points are individual samples. Dashed vertical line is the limit of detection. OTUs ordered alphabetically. * indicates that the OTU was unclassified at lower classification rank.

## References

1. Ducarmon QR, Zwittink RD, Hornung BVH, Schaik W van, Young VB, Kuijper EJ. 2019. Gut microbiota and colonization resistance against bacterial enteric infection. Microbiology and Molecular Biology Reviews 83. doi:10.1128/mmbr.00007-19.

2. Vardakas KZ, Trigkidis KK, Boukouvala E, Falagas ME. 2016. *Clostridium difficile* infection following systemic antibiotic administration in randomised controlled trials: A systematic review and meta-analysis. International Journal of Antimicrobial Agents 48:1–10. doi:10.1016/j.ijantimicag.2016.03.008.

3. Cammarota G, Ianiro G, Gasbarrini A. 2014. Fecal microbiota transplantation for the treatment of *Clostridium difficile* infection. Journal of Clinical Gastroenterology 48:693–702. doi:10.1097/mcg.0000000000000046.

4. Beran A, Sharma S, Ghazaleh S, Lee-Smith W, Aziz M, Kamal F, Acharya A, Adler DG. 2022. Predictors of fecal microbiota transplant failure in *Clostridioides difficile* infection. Journal of Clinical Gastroenterology Publish Ahead of Print. doi:10.1097/mcg.0000000000001667.

5. DeFilipp Z, Bloom PP, Soto MT, Mansour MK, Sater MRA, Huntley MH, Turbett S, Chung RT, Chen Y-B, Hohmann EL. 2019. Drug-resistant e. Coli bacteremia transmitted by fecal microbiota transplant. New England Journal of Medicine 381:2043–2050. doi:10.1056/nejmoa1910437.

6. Tvede M, Rask-Madsen J. 1989. Bacteriotherapy for chronic relapsing *Clostridium difficile* diarrhoea in six patients. The Lancet 333:1156–1160. doi:10.1016/s0140-6736(89)92749-9.

7. Lawley TD, Clare S, Walker AW, Stares MD, Connor TR, Raisen C, Goulding D, Rad R, Schreiber F, Brandt C, Deakin LJ, Pickard DJ, Duncan SH, Flint HJ, Clark TG, Parkhill J, Dougan G. 2012. Targeted restoration of the intestinal microbiota with a simple, defined bacteriotherapy resolves relapsing *Clostridium difficile* disease in mice. PLoS Pathogens 8:e1002995. doi:10.1371/journal.ppat.1002995.

8. Petrof EO, Gloor GB, Vanner SJ, Weese SJ, Carter D, Daigneault MC, Brown EM, Schroeter K, Allen-Vercoe E. 2013. Stool substitute transplant therapy for the eradication of *Clostridium difficile* infection: “RePOOPulating” the gut. Microbiome 1. doi:10.1186/2049-2618-1-3.

9. Kao D, Wong K, Franz R, Cochrane K, Sherriff K, Chui L, Lloyd C, Roach B, Bai AD, Petrof EO, Allen-Vercoe E. 2021. The effect of a microbial ecosystem therapeutic (MET-2) on recurrent *Clostridioides difficile* infection: A phase 1, open-label, single-group trial. The Lancet Gastroenterology & Hepatology 6:282–291. doi:10.1016/s2468-1253(21)00007-8.

10. Khanna S, Pardi DS, Kelly CR, Kraft CS, Dhere T, Henn MR, Lombardo M-J, Vulic M, Ohsumi T, Winkler J, Pindar C, McGovern BH, Pomerantz RJ, Aunins JG, Cook DN, Hohmann EL. 2016. A novel microbiome therapeutic increases gut microbial diversity and prevents recurrent *Clostridium difficile* infection. Journal of Infectious Diseases 214:173–181. doi:10.1093/infdis/jiv766.

11. McGovern BH, Ford CB, Henn MR, Pardi DS, Khanna S, Hohmann EL, O’Brien EJ, Desjardins CA, Bernardo P, Wortman JR, Lombardo M-J, Litcofsky KD, Winkler JA, McChalicher CWJ, Li SS, Tomlinson AD, Nandakumar M, Cook DN, Pomerantz RJ, Auninš JG, Trucksis M. 2020. SER-109, an investigational microbiome drug to reduce recurrence after *Clostridioides difficile* infection: Lessons learned from a phase 2 trial. Clinical Infectious Diseases 72:2132–2140. doi:10.1093/cid/ciaa387.

12. Taur Y, Coyte K, Schluter J, Robilotti E, Figueroa C, Gjonbalaj M, Littmann ER, Ling L, Miller L, Gyaltshen Y, Fontana E, Morjaria S, Gyurkocza B, Perales M-A, Castro-Malaspina H, Tamari R, Ponce D, Koehne G, Barker J, Jakubowski A, Papadopoulos E, Dahi P, Sauter C, Shaffer B, Young JW, Peled J, Meagher RC, Jenq RR, Brink MRM van den, Giralt SA, Pamer EG, Xavier JB. 2018. Reconstitution of the gut microbiota of antibiotic-treated patients by autologous fecal microbiota transplant. Science Translational Medicine 10. doi:10.1126/scitranslmed.aap9489.

13. Reigadas E, Prehn J van, Falcone M, Fitzpatrick F, Vehreschild MJGT, Kuijper EJ, Bouza E. 2021. How to: Prophylactic interventions for prevention of *Clostridioides difficile* infection. Clinical Microbiology and Infection 27:1777–1783. doi:10.1016/j.cmi.2021.06.037.

14. Lesniak NA, Schubert AM, Sinani H, Schloss PD. 2021. Clearance of *Clostridioides difficile* colonization is associated with antibiotic-specific bacterial changes. mSphere 6. doi:10.1128/msphere.01238-20.

15. Ghimire S, Roy C, Wongkuna S, Antony L, Maji A, Keena MC, Foley A, Scaria J. 2020. Identification of *Clostridioides difficile-inhibiting* gut commensals using culturomics, phenotyping, and combinatorial community assembly. mSystems 5. doi:10.1128/msystems.00620-19.

16. Auchtung JM, Preisner EC, Collins J, Lerma AI, Britton RA. 2020. Identification of simplified microbial communities that inhibit *Clostridioides difficile* infection through dilution/extinction. mSphere 5. doi:10.1128/msphere.00387-20.

17. Tomkovich S, Stough JMA, Bishop L, Schloss PD. 2020. The initial gut microbiota and response to antibiotic perturbation influence *Clostridioides difficile* clearance in mice. mSphere 5. doi:10.1128/msphere.00869-20.

18. Tomkovich S, Taylor A, King J, Colovas J, Bishop L, McBride K, Royzenblat S, Lesniak NA, Bergin IL, Schloss PD. 2021. An osmotic laxative renders mice susceptible to prolonged *Clostridioides difficile* colonization and hinders clearance. mSphere 6. doi:10.1128/msphere.00629-21.

19. Buffie CG, Bucci V, Stein RR, McKenney PT, Ling L, Gobourne A, No D, Liu H, Kinnebrew M, Viale A, Littmann E, Brink MRM van den, Jenq RR, Taur Y, Sander C, Cross JR, Toussaint NC, Xavier JB, Pamer EG. 2014. Precision microbiome reconstitution restores bile acid mediated resistance to *Clostridium difficile*. Nature 517:205–208. doi:10.1038/nature13828.

20. Reeves AE, Koenigsknecht MJ, Bergin IL, Young VB. 2012. Suppression of *Clostridium difficile* in the gastrointestinal tracts of germfree mice inoculated with a murine isolate from the family Lachnospiraceae. Infection and Immunity 80:3786–3794. doi:10.1128/iai.00647-12.

21. Leslie JL, Vendrov KC, Jenior ML, Young VB. 2019. The gut microbiota is associated with clearance of *Clostridium difficile* infection independent of adaptive immunity. mSphere 4. doi:10.1128/mspheredirect.00698-18.

22. Nagao-Kitamoto H, Leslie JL, Kitamoto S, Jin C, Thomsson KA, Gillilland MG, Kuffa P, Goto Y, Jenq RR, Ishii C, Hirayama A, Seekatz AM, Martens EC, Eaton KA, Kao JY, Fukuda S, Higgins PDR, Karlsson NG, Young VB, Kamada N. 2020. Interleukin-22-mediated host glycosylation prevents *Clostridioides difficile* infection by modulating the metabolic activity of the gut microbiota. Nature Medicine 26:608–617. doi:10.1038/s41591-020-0764-0.

23. Byndloss MX, Olsan EE, Rivera-Chávez F, Tiffany CR, Cevallos SA, Lokken KL, Torres TP, Byndloss AJ, Faber F, Gao Y, Litvak Y, Lopez CA, Xu G, Napoli E, Giulivi C, Tsolis RM, Revzin A, Lebrilla CB, Bäumler AJ. 2017. Microbiota-activated PPAR-signaling inhibits dysbiotic Enterobacteriaceae expansion. Science 357:570–575. doi:10.1126/science.aam9949.

24. Winter SE, Lopez CA, Bäumler AJ. 2013. The dynamics of gut-associated microbial communities during inflammation. EMBO reports 14:319–327. doi:10.1038/embor.2013.27.

25. Lesniak NA, Schubert AM, Flynn KJ, Leslie JL, Sinani H, Bergin IL, Young VB, Schloss PD. 2022. The gut bacterial community potentiates Clostridioides difficile infection severity. doi:10.1101/2022.01.31.478599.

26. Nakashima T, Fujii K, Seki T, Aoyama M, Azuma A, Kawasome H. 2021. Novel gut microbiota modulator, which markedly increases *Akkermansia muciniphila* occupancy, ameliorates experimental colitis in rats. Digestive Diseases and Sciences. doi:10.1007/s10620-021-07131-x.

27. Stein RR, Bucci V, Toussaint NC, Buffie CG, Rätsch G, Pamer EG, Sander C, Xavier JB. 2013. Ecological modeling from time-series inference: Insight into dynamics and stability of intestinal microbiota. PLoS Computational Biology 9:e1003388. doi:10.1371/journal.pcbi.1003388.

28. Flynn KJ, Ruffin MT, Turgeon DK, Schloss PD. 2018. Spatial variation of the native colon microbiota in healthy adults. Cancer Prevention Research 11:393–402. doi:10.1158/1940-6207.capr-17-0370.

29. Guilloux C-A, Lamoureux C, Beauruelle C, Héry-Arnaud G. 2021. Porphyromonas: A neglected potential key genus in human microbiomes. Anaerobe 68:102230. doi:10.1016/j.anaerobe.2020.102230.

30. Seekatz AM, Theriot CM, Rao K, Chang Y-M, Freeman AE, Kao JY, Young VB. 2018. Restoration of short chain fatty acid and bile acid metabolism following fecal microbiota transplantation in patients with recurrent *Clostridium difficile* infection. Anaerobe 53:64–73. doi:10.1016/j.anaerobe.2018.04.001.

31. Rinttila T, Kassinen A, Malinen E, Krogius L, Palva A. 2004. Development of an extensive set of 16S rDNA-targeted primers for quantification of pathogenic and indigenous bacteria in faecal samples by real-time PCR. Journal of Applied Microbiology 97:1166–1177. doi:10.1111/j.1365-2672.2004.02409.x.

32. Sorg JA, Dineen SS. 2009. Laboratory maintenance of *Clostridium difficile*. Current Protocols in Microbiology 12. doi:10.1002/9780471729259.mc09a01s12.

33. Winston JA, Thanissery R, Montgomery SA, Theriot CM. 2016. Cefoperazone-treated mouse model of clinically-relevant Δ*Clostridium difficile* strain r20291. Journal of Visualized Experiments. doi:10.3791/54850.

34. Kozich JJ, Westcott SL, Baxter NT, Highlander SK, Schloss PD. 2013. Development of a dual-index sequencing strategy and curation pipeline for analyzing amplicon sequence data on the MiSeq illumina sequencing platform. Applied and Environmental Microbiology 79:5112–5120. doi:10.1128/aem.01043-13.

35. Schloss PD, Westcott SL, Ryabin T, Hall JR, Hartmann M, Hollister EB, Lesniewski RA, Oakley BB, Parks DH, Robinson CJ, Sahl JW, Stres B, Thallinger GG, Horn DJV, Weber CF. 2009. Introducing mothur: Open-source, platform-independent, community-supported software for describing and comparing microbial communities. Applied and Environmental Microbiology 75:7537–7541. doi:10.1128/aem.01541-09.

36. Wang Q, Garrity GM, Tiedje JM, Cole JR. 2007. Naïve bayesian classifier for rapid assignment of rRNA sequences into the new bacterial taxonomy. Applied and Environmental Microbiology 73:5261–5267. doi:10.1128/aem.00062-07.

37. Yue JC, Clayton MK. 2005. A similarity measure based on species proportions. Communications in Statistics - Theory and Methods 34:2123–2131. doi:10.1080/sta-200066418.

38. Benjamini Y, Hochberg Y. 1995. Controlling the false discovery rate: A practical and powerful approach to multiple testing. Journal of the Royal Statistical Society: Series B (Methodological) 57:289–300. doi:10.1111/j.2517-6161.1995.tb02031.x.

39. Kurtz ZD, Müller CL, Miraldi ER, Littman DR, Blaser MJ, Bonneau RA. 2015. Sparse and compositionally robust inference of microbial ecological networks. PLOS Computational Biology 11:e1004226. doi:10.1371/journal.pcbi.1004226.

